# TAD cliques predict key features of chromatin organization

**DOI:** 10.1101/2020.11.01.363903

**Authors:** Tharvesh M. Liyakat Ali, Annaël Brunet, Philippe Collas, Jonas Paulsen

**Affiliations:** Department of Molecular Medicine, Institute of Basic Medical Sciences, Faculty of Medicine, University of Oslo, Oslo, Norway; Department of Immunology and Transfusion Medicine, Oslo University Hospital, Oslo, Norway; Institute of Biosciences, Faculty of Mathematics and Natural Sciences, University of Oslo, Oslo, Norway

**Keywords:** 3D genome, chromatin conformation, Hi-C, TAD, CTCF motif

## Abstract

Processes underlying genome 3D organization and domain formation in the mammalian nucleus are not completely understood. Multiple processes such as transcriptional compartmentalization, DNA loop extrusion and interactions with the nuclear lamina dynamically act on chromatin at multiple levels. Here, we explore long-range interaction patterns between topologically associated domains (TADs) in several cell types. We find that this is connected to many key features of chromatin organization, including open and closed compartments, compaction and loop extrusion processes. We find that domains that form large TAD cliques tend to be repressive across cell types, when comparing gene expression, LINE/SINE repeat content and chromatin subcompartments. Further, TADs in large cliques are larger in genomic size, less dense and depleted of convergent CTCF motifs, in contrast to smaller and denser TADs formed by a loop extrusion processes. Our results shed light on the organizational principles that govern repressive and active domains in the human genome.

## Introduction

Spatial organization and packaging of the genome are important for proper regulation of gene expression and are often altered in disease [1]. Understanding the underlying organizational principles of 3D genome architecture requires a multi-scale and multi-scope approach. At higher-order levels, chromosomes seem to organize into two large A and B compartments which can be computed from the first eigenvector of a principal component analysis (PCA) of a correlation Hi-C matrix at low resolution (e.g. 1 megabase [Mb]) [2]. By definition, A compartments constitute open/active parts of the genome, while B compartments make up the remaining inactive parts. Increasing the resolution, and thus decreasing the bin size of a Hi-C matrix, reveals a finer delineation of compartments into subcompartments [3]. Zooming further on the diagonal of the Hi-C matrix reveals nested levels of high-frequency interactions delineated by relatively abrupt boundaries between them, referred to as topologically-associated domains (TADs) [4, 5]. Several processes together likely shape the chromosomal interaction patterns observed in Hi-C matrices. It has notably been proposed that a phase-separation process could explain the formation of heterochromatin compartments [6, 7], and a loop-extrusion model could explain TAD formation and dynamics [8, 9]. For most genomic regions, multiple processes act simultaneously within and between cells in a population to spatially organize the genome at multiple levels [10, 11].

Based on analysis of the *Drosophila* genome, high-resolution Hi-C data show that compartments of very small sizes can be computed from an eigenvector analysis similar to what has previously been applied on low-resolution Hi-C data [12]. These compartments, termed compartment domains, correspond almost perfectly to transcription state transitions in the *Drosophila* genome [12]. Such compartment domains are also found in mammalian genomes [12]. However in addition, chromatin looping events involving CCCTC-binding factor (CTCF) seem to play a prominent role in the formation of TADs [3], in particular through loop extrusion processes [8, 9]. Computational simulations reveal that small compartment domains are partially suppressed by loop extrusion processes counteracting their segregation [10]. The view of mammalian 3D genome organization is thus becoming increasingly complex, and further classification of the different types of chromatin domains has been suggested [13]. We have recently shown that long-range TAD-TAD interactions can occur in the form of TAD cliques, which we have defined as an assembly of at ≥ 3 TADs that are fully connected pairwise in Hi-C data [14]. TAD cliques associate with key organizational processes during adipose stem cell differentiation, notably by stabilizing heterochromatin at the nuclear periphery, through lamina-associated domains (LADs) [14]. Here, we explore the properties of TADs engaging in TAD-TAD interactions in four human cell lines. We find that TADs that belong to large or small cliques display distinct genomic features. Most significantly, TADs in large cliques are depleted of convergent CTCF-motifs at their boundaries, unlike ‘classical’ TADs explained by chromatin loop extrusion processes. Our findings shed further light on long-range TAD-TAD interactions and indicate that they constitute an important structural feature of the genome.

## Results

Long-range interactions between linearly non-contiguous TADs, together with interactions between TADs and the nuclear lamina via LADs, shape genome architecture during differentiation of adipose stem cells [14]. To further explore such TAD-TAD interactions in other cell types, we analyzed TADs in four human cell lines (HMEC, HUVEC, IMR90, K562) for which high-resolution Hi-C and gene expression information is available [15] (see **Table S1** for accession numbers). Using Armatus [16] (see Methods), we identified a total of 5502-6008 TADs in each cell line (**Table S2**), consistent with our previous findings in primary adipose stem cells using the same algorithm [14]. These TADs display similar characteristics as shown earlier [4, 5, 14], with marked boundary structures and sizes in the range ∼0.2-1 Mb (**Fig. 1A**).

**Fig. 1.**
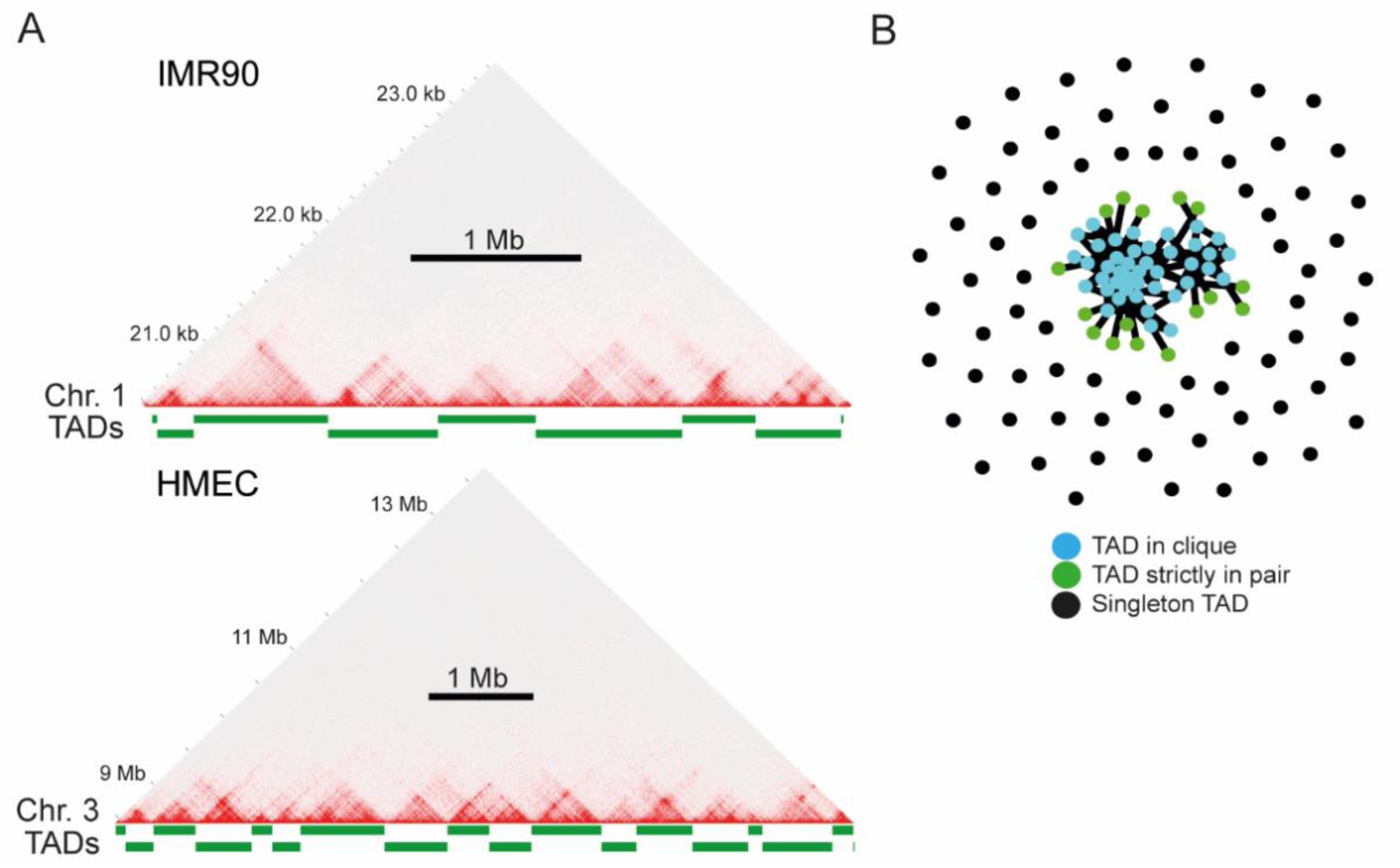
TADs and TAD interaction networks. (A) Examples of TADs identified in Hi-C matrices of IMR90 and HMEC cells. Delineation of Armatus TADs is shown as green bars. (B) TAD networks: graph representation of TADs in clique, binary interacting TADs (TADs in pairs only) and singleton TADs for chromosome 18 in IMR90 cells.

### TAD-TAD interactions, TAD cliques and gene repression

To determine the presence of TAD-TAD interactions from the Hi-C data in HMEC, HUVEC, IMR90 and K562 cells, we used the Non-central Hypergeometric model as done previously [14, 17, 18]. We find a total of ∼6000-8300 significant intra-chromosomal interactions (IMR90: 8300; HMEC: 7309; HUVEC: 5934; K562: 7823). Interactions between TADs are configured as complex networks of strictly pairwise interactions, or involving multiple interactions, with enrichments and depletions of contacts across chromosomes, as exemplified for chromosome 18 in IMR90 cells (**Fig. 1B**).

TADs can engage in interactions with multiple TADs, some forming cliques (where all TADs interact pairwise [14]), some not. In addition, a TAD can be part of one or more cliques of different size (the size of a clique is defined by the number of TADs that comprise it). We use the term of ‘TAD maximal clique size’, referring to the size of the largest clique a given TAD belongs to [14]. Maximal clique sizes were determined for all four cell types, as done previously using the Bron-Kerbosch algorithm [14]. We find that across cell lines, 1189-1554 TADs engage in associations with at least two other linearly non-contiguous TADs, forming cliques of size ≥ 3 (**Fig. 2A**; **Table S2**). This represents 21-27% of all TADs in these cell lines (**Fig. 2A**), supporting the view that TAD cliques constitute a significant feature of higher-order genome topology. As previously reported [14], genes residing within TADs in cliques are expressed at a lower level than those in TADs outside cliques (**Fig. 2B**), corroborating the repressive nature of TAD cliques.

**Fig. 2.**
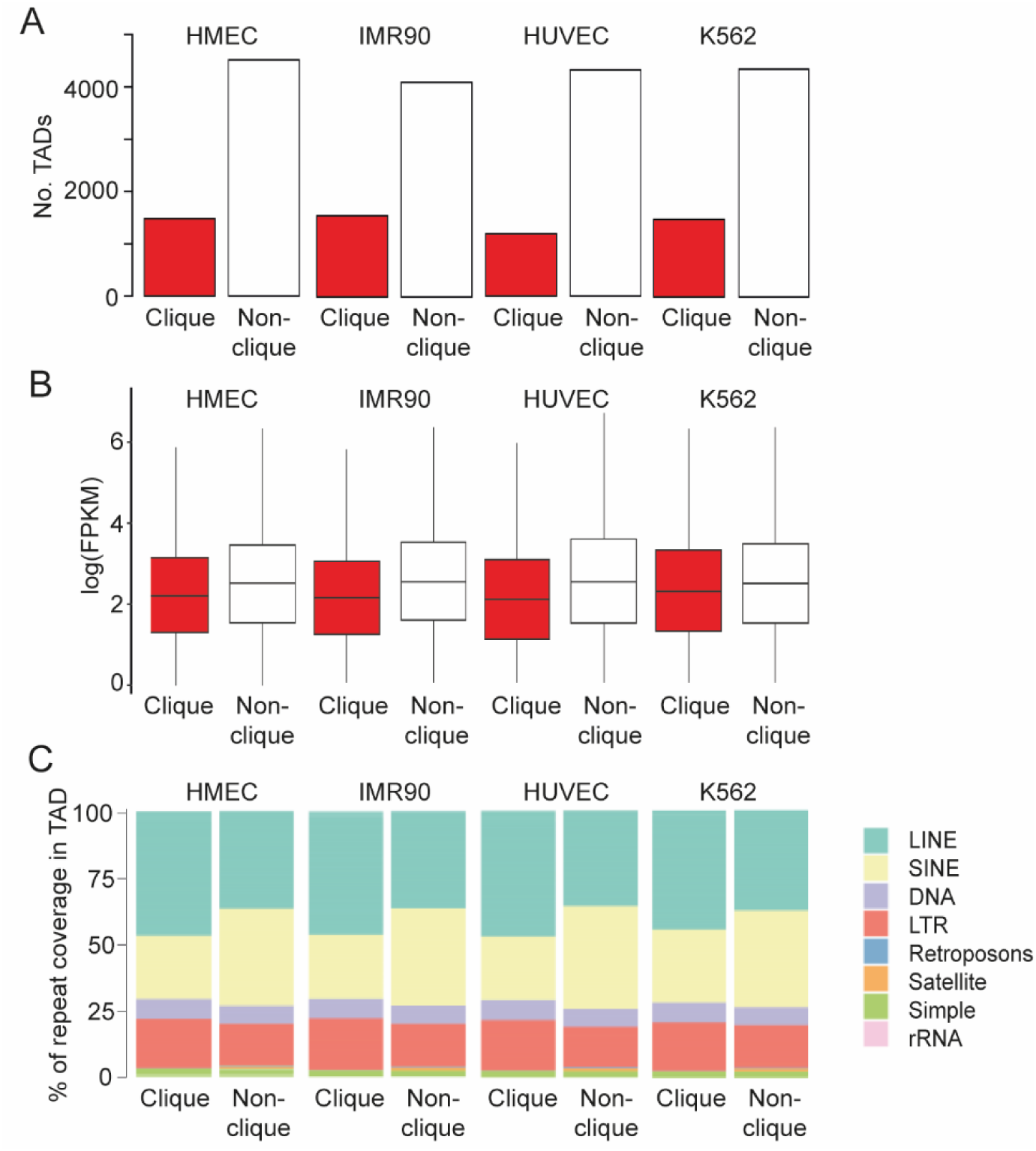
Genomic characterization of TADs in cliques. (A) Number of TADs (Armatus) in cliques and outside cliques in indicated cell types, identified from publicly available Hi-C data. (B) Distribution of gene expression levels in TADs in cliques and outside cliques. (C) Proportion of TAD coverage by indicated repeat classes in cliques and outside cliques.

Retrotransposons play an increasingly appreciated role in gene expression and chromatin structure regulation [19, 20]. Relevant for genome architecture is evidence that long interspersed elements (LINEs) and short interspersed elements (SINEs) can modulate transcription by altering chromatin composition [21] and structure [22]. Notably, LINEs and SINEs may act as euchromatin-heterochromatin boundary elements confining gene expression to the proper compartment [22] or play a role in the formation of silent domains [23]. The relationship between retrotransposons and long-range TAD-TAD interactions has however not been thoroughly examined. We thus investigated the genomic distribution of repeat classes across TADs in and outside cliques. We find a systematic enrichment of LINE coverage, and correspondingly a depletion of SINE coverage, for TADs in cliques compared to TADs outside cliques (**Fig 2C**). Other repeat classes show limited if any differential coverage (**Fig. 2C**). As LINE elements are implicated in heterochromatin formation [23], this finding further establishes TAD cliques as repressive sub-compartments of the genome.

### Genomic characterization of TADs in cliques

As TADs usually are defined solely from short-range Hi-C contact enrichments separated by sharp boundaries [4, 5], the processes underlying their formation could vary between different TADs. Several partially independent processes have been proposed to shape TADs [11, 13]. Loop extrusion has been proposed as an underlying process in TAD formation [8, 9], whereas phase separation has been suggested as a mechanism of compartmentalization of chromatin [6, 7]. In the human genome, a combination of these processes seems to underline the delineation of many TADs [12].

Visualization of Hi-C contact patterns within TADs in cliques reveals a distinct contact feature often characterized by larger and less densely interacting domains compared to TADs not in cliques (exemplified in **Fig. 3A**). To investigate this further, we determined the distribution of TAD sizes for TADs identified as singletons, TADs interacting only in pairs (binary interacting TADs), and TADs belonging to cliques of increasing sizes. At the whole genome level, we note a linear relationship between clique size and median size of TADs in these cliques (**Fig. 3B**). Further, genome-wide analysis of Hi-C contact densities within TADs in varying TAD clique classes indicates that TADs in larger cliques systematically display a less dense contact pattern than singleton and binary interacting TADs (**Fig. 3C**).

**Fig. 3.**
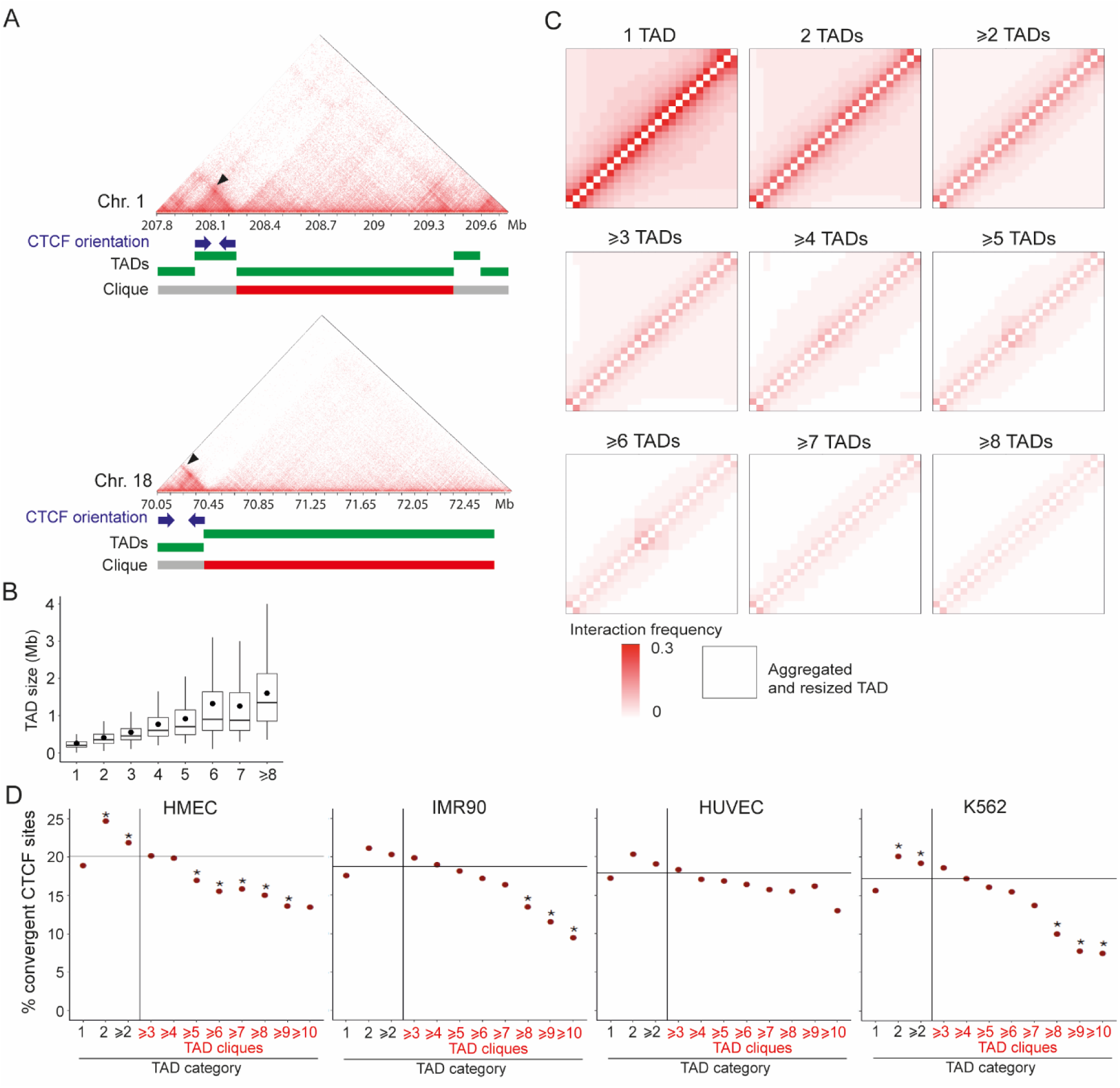
TADs in cliques display less dense interaction patterns than singleton or binary interacting TADs and are impoverished in convergent CTCF motifs. (A) Hi-C matrices for segments of chromosomes 1 and 18 (IMR90 cells); Armatus TADs are delineated by green bars. A TAD belonging to a clique is indicated by a red bar (gray otherwise). Small TADs containing dense chromosomal interactions display convergent CTCF motifs at their boundaries (blue arrows); arrowheads, corner interaction peak. (B) TAD size distribution in IMR90 cells as a function of clique size (3 to ≥ 8) the TADs belong to. Bar, median; dot, mean. (C) Mean interaction frequencies in aggregated and resized TADs. Each matrix is for aggregated TADs in the indicated categories. (D) Percentage of convergent CTCF motifs at the boundaries of TADs categorized as shown. The horizontal bar represents the average percentage of convergent CTCF motifs in all TADs genome-wide. *Binomial test; see **Table S3** for statistics.

The presence and orientation of CTCF motifs at each TAD boundary has been shown to be indicative of TAD formation and stability [3, 24]. Given our previous observation of higher density interactions within small TADs than in large TADs, we explored the enrichment of convergent CTCF motifs at the boundaries of TADs in the cell lines examined in our study. Interestingly, convergent CTCF motifs and corner peaks seem less prominent for TADs in cliques than for TADs not in cliques (**Fig. 3A**, blue arrows and black arrowheads). We therefore hypothesized that the process shaping TADs in cliques might be distinct from that shaping TADs outside cliques.

To test this hypothesis, we computed genome-wide enrichment scores of convergent CTCF motifs for (i) singleton TADs, (ii) TADs involved in binary interactions and (iii) TADs in cliques of increasing size (**Fig. 3D**). We find that TADs engaging in interaction with only one other TAD are the most enriched in convergent CTCF motifs at their boundaries, whereas TAD in cliques of increasing size show a gradual decrease in convergent CTCF motif enrichment (**Fig. 3D**). In fact, in large cliques (≥ 5 TADs), convergent CTCF motifs are depleted compared to the average convergent CTCF motif enrichment across all TADs in the genome. For cliques of ≥ 5-8 TADs in HMEC, IMR90 and K562 cells, this depletion is statistically significant (**Table S3**). Singleton TADs are less enriched in convergent CTCF motifs than binary interacting TADs, and also depleted compared to the genome-wide average (**Fig. 3D**). These trends are systematic across the four cell lines, suggesting a general relationship. Since convergent CTCF motifs are implicated in loop extrusion processes, our data suggest that binary interacting TADs are more likely to form by loop extrusion compared to TADs in cliques and, to a lesser extent, singleton TADs. The implication from our findings that TADs formed by loop extrusion are more likely to engage in binary interactions may reflect the previously characterized nested structures of these TADs [13, 25].

### Relationship between TAD-cliques and compartments

Eigenvector analysis of high-resolution Hi-C data has previously been used to determine regions with a genomic size similar to TADs that segregate into six different subcompartments [3]. These have been shown to correspond to distinct types of active (subcompartment A1 and A2) and inactive (subcompartments B1-B4) regions of the genome [3]. The clique pattern of TAD-TAD interactions suggests a relationship with these subcompartments: we hypothesized that TADs in cliques behave as small, individual compartments, possibly suggesting localized compartmentalization as a separate mechanism of TAD formation. To examine this possibility, we determined the overlap of subcompartment segments to TADs in cliques. Using the Jaccard index (JI) as a measure of the relative overlap between each TAD and its overlapping subcompartment(s), we found only a limited correspondence between these (median JI 0.1-0.3), irrespective of subcompartment type and cell type (**Fig. 4A**). Notwithstanding, for all cell types except K562, A1 subcompartment overlap diminishes as TAD clique size increases (**Fig. 4A**). For all cell types, overlap with B2 and B3 subcompartments tend to increase for larger clique sizes (**Fig. 4A**). We conclude from these observations that TAD cliques are distinct from previously annotated subcompartments.

**Fig. 4.**
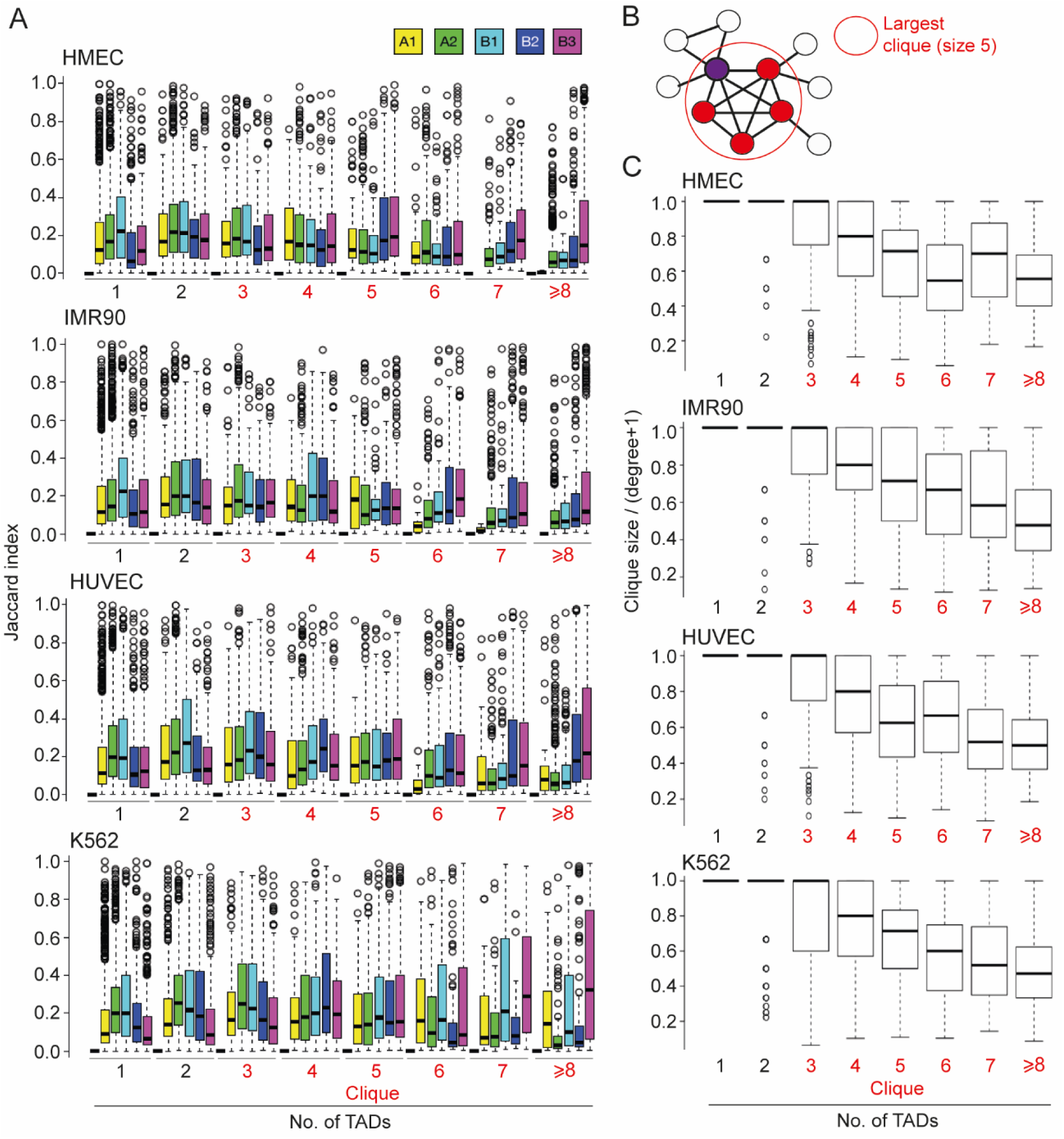
TADs in large cliques interact with a large number of TADs also outside the clique. (A) Overlap between singleton TADs, binary interacting TADs and TADs in cliques (of indicated size) with A and B compartment subtypes. (B) Concept of ‘degree’ of TAD interactions. A given TAD (purple node) can belong to a clique of, here, size 3 (containing two other TADs [white nodes]) and a clique of size 5 (red nodes); the latter is the ‘maximal clique size’ (see main text). The total number, or ‘degree’, of interactions the purple TADs engages in is 7 and are materialized by 7 edges. In this example, the ratio of (clique size / (degree+1)) is 5/(7+1) = 0.625 (see panel C). (C) Ratios of (clique size / (degree+1)) for TADs identified as singletons, binary interacting and in cliques. The graphs consistently show that larger the clique size, the lower the ratio, i.e. the greater the number of inter-TAD interactions a TAD engages in *outside* the clique. Note that for singleton TADs, this ratio is (trivially) always 1.

To further understand the interaction patterns of TADs, we explored the relationship between TAD-TAD interactions and clique size, as this could shed light on whether TAD cliques might constitute an exclusive mode of regionalization of the genome rather than highly interacting compartments. More explicitly, we examined the relationship between the total number of TADs a given TAD interacts with and the size of the largest clique this TAD belongs to (**Fig. 4B**). **Figure 4C** shows the ratio of (largest) clique size to the total number (‘degree’) of interactions of each TAD, for increasing clique sizes; this reflects how many of each TAD’s interactions are accounted for by their interactions in cliques. Consistently across cell types, we find that larger cliques tend to interact with a greater number of other TADs also outside of the clique (resulting in lower clique size / interaction degree ratios; **Fig. 4C**). We speculate that this may result from heterochromatin being more compact and interacting more closely with other heterochromatin regions, further supporting a view of preferred homotypic chromatin associations [8, 24, 26]. In contrast, the lower density of inter-TAD interaction, manifested by high ratios involving TAD singletons or binary interacting TADs or small cliques (**Fig. 5C**) reflects more open chromatin configurations which are less interactive, except within TADs or with neighboring TADs (see e.g. **Fig. 3A**).

## Discussion

We report a genomic assessment of TADs in cliques, large ‘multi-TAD’ assemblies detected from ensemble Hi-C data. Our results suggest that a subset of TADs serves regulatory function through the formation of long-range interactions, yet the definition of TADs has recently been challenged [11]. We also note that the nature of Hi-C contact domains is not fully understood. For example, Rowley et al. [27] report that 1939 (23%) TAD boundaries cannot be explained by neither extrusion nor compartmental processes. The TADs in cliques reported here are characterized by being larger and less dense than typical TADs, and with a depletion of convergent CTCF motifs at their boundaries. This clearly suggests that chromatin loop extrusion cannot explain the formation of these TADs. Due to their large size, TADs in large cliques also do not fit the definition of compartment domains, which are typically smaller than TADs [12, 27]. The question remains therefore of which processes shape these domains. Their previously reported association with the nuclear lamina [14], and their association with repressive chromatin marks, suggest that heterochromatin tethering protein factors such as CBX5/HP1α [28] could be involved. Knockdown of these factors in combination with Hi-C analysis and TAD clique identification could therefore elucidate this further.

In a recently suggested classification of Hi-C domains, TADs in TAD cliques would probably be classified as type 3 ‘Compartment domain only: un-nested no-corner-dot compartment domain’ [13]. The large genomic size and relatively lower interaction density of these TADs compared to previously described compartment domains could however be indicative of a separate formation process.

We have relied on the Armatus TAD caller [16] for the delineation of TADs. This choice was based on testing a range of TAD callers and selecting the one that provided the most reproducible and visually pronounced TADs. It is however inevitable that some of the called TADs may be less well-defined using this algorithm. Even if we have taken a TAD-based approach, our findings do not rule out that compartment domains not identified as TAD cliques serve important regulatory functions.

We find that binary interacting TADs, unlike singleton TADs, are the most enriched in convergent CTCF motifs. The explanation for this could be that binary interacting TADs are indicative of a nested TAD structure. These nested TAD structures have been shown to often be found for domains caused by loop-extrusion processes [13].

We find TAD-cliques across different cell types, suggesting that TAD cliques are general phenomena not only linked to cell differentiation. In this regard, TAD cliques constitute an interesting and important chromatin feature for further study, since they link local interaction patterns (i.e. TADs and compartment domains) to higher order organization (i.e. compartments and LADs). A deeper characterization of TAD cliques across cell and tissue types might further elucidate these relationships. Also, single-cell analysis, including high-throughput imaging, might reveal whether TAD cliques result from an aggregation of interactions across cells, or exist within single cells. Taken together, our results shed further light on the increasingly complex picture of multiscale chromatin organization.

## Methods

### Hi-C data

To uniformly process all Hi-C data used in this study, raw data were downloaded from ENCODE [15] and processed using the HiC-Pro pipeline [29] (https://github.com/nservant/HiC-Pro). First, the paired-end sequences were mapped to the hg38 reference genome using Bowtie2 [30] with default parameters preset in HiC-Pro configuration file. Unmapped, multi-mapped, singletons and low map quality reads were removed and only uniquely mapped reads were used for binning, normalizing and generating Hi-C matrices. The pipeline produced raw and normalized interaction frequency matrices. For further analyses, 5 kb and 50 kb resolution raw matrices were used for all cell lines. We used the hicpro2juicebox.sh script from HiC-Pro to convert matrices into .hic files for visualization with Juicebox [31] (https://github.com/theaidenlab/juicebox).

### TAD calling

TADs were called using Armatus v2.1.0 [16] (https://github.com/kingsfordgroup/armatus) using a gamma of 1.2 for all cell lines. Genomic regions not defined as TADs by Armatus were nevertheless included to ensure full genome segmentation. TADs were visualized using Juicebox (**Fig. 1A**).

### Identification of TAD-TAD interactions

TAD-TAD interactions were identified using the NCHG (Non-central Hypergeometric model) tool [17]. Hi-C contacts were aggregated to generate TAD-TAD interaction matrices for each cell line. NCHG was used to calculate P-values for each TAD pair. Then, we performed multiple testing correction with a false discovery rate (FDR) < 1% using the Benjamini-Hochberg method. The resulting significant interactions were filtered by requiring a five-fold enrichment of observed over expected contacts based on genomic distance.

The network configuration of TAD-TAD interactions (**Fig. 1B**) was generated using the igraph R package [32] (https://github.com/igraph/rigraph). The igraph layout was made using the 131 TADs identified in chromosome 18 of IMR90 cells. We used the ‘graphopt’ algorithm setting the charge parameter to 0.03 while the remaining parameters were left as default. Each node was colored-coded based on the degree of interactions.

### TAD clique calling

As we reported earlier [14], significant TAD-TAD interactions were represented as a graph using the NetworkX Python library (http://networkx.github.io/). In the graph, TADs are represented by nodes and significant interactions between them are represented by edges. Maximal TAD clique sizes were calculated using the Bron-Kerbosch algorithm [33]. Maximal clique size (*k*) was assigned to each TAD, where *k* is the size of the largest TAD clique to which the TAD belongs to.

### Repeat analysis

The repeat mask file for the hg38 genome assembly was downloaded from the UCSC genome browser[34] (http://hgdownload.cse.ucsc.edu/goldenpath/hg38/database/rmsk.txt.gz). From the repeat mask file, the following repeats were selected for further analysis: LINE, SINE, LTR, retrotransposons, rRNA, satellite, simple and DNA. The repeat contents for each TAD were calculated using the bedtools coverage option [35] and plots generated using the ggplot2 R package.

### Aggregated TADs

Intra-TAD interaction frequencies for each TAD in IMR90 cells at 5 kb resolution was extracted from the Hi-C matrix. As the genomic length of TADs differs, so do the sizes of intra-TAD interaction frequency matrices. Therefore, all TADs were resized to a 25 x 25 matrix using the ‘nearest’ algorithm from the OpenImageR R package (https://github.com/mlampros/OpenImageR). The element-wise mean was calculated for all TADs of a given category (based on clique size) to produce the mean matrix for that category.

### CTCF motif orientation analysis

Processed CTCF peak files in NarrowPeak format for all cell lines were downloaded from ENCODE [15]. The GimmeMotif [36], a transcription factor analysis tool, was used to call all motifs from the peak files using the ‘scan’ option passing the ‘JASPAR2020_vertebrates’ PFM file. From the resulting bed file, CTCF peaks were extracted with information on the orientation of CTCF binding. Python and R scripts were used to calculate the CTCF orientations at TAD boundaries.

### Scripting

All scripts for data analyses in this study were written using R, Python and Bash. The scripts can be found on GitHub (https://github.com/tharvesh/paper3).

## Acknowledgements

This work was supported by the Research Council of Norway (PC), the Norwegian Cancer Society (PC) and the University of Oslo (PC, JP).

## Author contributions

TMLA, PC and JP wrote the manuscript. AB, PC and JP supervised the work. All authors approved the final version.

## Competing interests

The authors declare that they have no competing interests.

## List of abbreviations

ChIP: chromatin immunoprecipitation;
JI: Jaccard index;
LAD: lamina-associated domain;
PCA: principal component analysis;
TAD: topologically-associated domain

## Supplementary Tables

**Table S1.** ENCODE data accession numbers

| Data type | IMR90 | HMEC | HUVEC | K562 |
| --- | --- | --- | --- | --- |
| Hi-C raw data | SRR1658672 | SRR1658680 | SRR1658709 | SRR1658693 |
|  | SRR1658673 | SRR1658681 | SRR1658710 | SRR1658694 |
|  | SRR1658674 | SRR1658682 | SRR1658711 | SRR1658695 |
|  | SRR1658675 | SRR1658683 | SRR1658712 | SRR1658696 |
|  | SRR1658676 | SRR1658684 | SRR1658713 | SRR1658697 |
|  | SRR1658677 | SRR1658685 | SRR1658714 | SRR1658698 |
|  | SRR1658678 |  |  | SRR1658699 |
|  |  |  |  | SRR1658700 |
|  |  |  |  | SRR1658701 |
|  |  |  |  | SRR1658702 |
| RNA-seq data | ENCFF244VME | ENCFF526QYF | ENCFF238WEU | ENCFF104VTJ |
| ChIP-seq data | ENCFF307XFM | ENCFF641CHY | ENCFF536OTV | ENCFF873LHT |

**Table S2.** Numbers of TADs in cliques and non-cliques

| Cell type | Number of TADs |  |  |
| --- | --- | --- | --- |
|  | Clique | Non-clique | Total |
| HMEC | 1486 | 4522 | 6008 |
| IMR90 | 1554 | 4095 | 5649 |
| HUVEC | 1189 | 4313 | 5502 |
| K562 | 1488 | 4332 | 5820 |

**Table S3.** Statistics on the depletion of convergent CTCF sites in TADs as a function of clique size

| Clique category | Confident1 | Confident2 | TADs with conv. CTCF motifs | No. TADs in category | % TADs with conv. CTCF in genome | P-value |
| --- | --- | --- | --- | --- | --- | --- |
| **IMR90** |  |  |  |  |  |  |
| ==2 | 0.1835 | 0.2405 | 173 | 820 | 0.1871 | 0.0809 |
| >=2 | 0.1866 | 0.2194 | 481 | 2374 | 0.1871 | 0.0547 |
| >=3 | 0.1786 | 0.2189 | 308 | 1554 | 0.1871 | 0.2688 |
| >=4 | 0.1669 | 0.2135 | 214 | 1130 | 0.1871 | 0.8488 |
| >=5 | 0.1560 | 0.2084 | 157 | 867 | 0.1871 | 0.6951 |
| >=6 | 0.1443 | 0.2021 | 118 | 687 | 0.1871 | 0.3278 |
| >=7 | 0.1321 | 0.1986 | 82 | 502 | 0.1871 | 0.1880 |
| >=8 | 0.1008 | 0.1743 | 48 | 357 | 0.1871 | 0.0098 |
| >=9 | 0.0756 | 0.1646 | 25 | 218 | 0.1871 | 0.0053 |
| >=10 | 0.0538 | 0.1508 | 15 | 159 | 0.1871 | 0.0021 |
| **HMEC** |  |  |  |  |  |  |
| ==2 | 0.2182 | 0.2757 | 220 | 894 | 0.2006 | 0.0010 |
| >=2 | 0.2016 | 0.2352 | 519 | 2380 | 0.2006 | 0.0358 |
| >=3 | 0.1811 | 0.2225 | 299 | 1486 | 0.2006 | 0.9484 |
| >=4 | 0.1744 | 0.2239 | 205 | 1034 | 0.2006 | 0.8766 |
| >=5 | 0.1433 | 0.1978 | 129 | 762 | 0.2006 | 0.0299 |
| >=6 | 0.1256 | 0.1878 | 85 | 549 | 0.2006 | 0.0065 |
| >=7 | 0.1218 | 0.1997 | 57 | 361 | 0.2006 | 0.0417 |
| >=8 | 0.1100 | 0.1982 | 41 | 273 | 0.2006 | 0.0409 |
| >=9 | 0.0904 | 0.1921 | 26 | 192 | 0.2006 | 0.0239 |
| >=10 | 0.0788 | 0.2091 | 16 | 119 | 0.2006 | 0.0850 |
| **HUVEC** |  |  |  |  |  |  |
| ==2 | 0.1749 | 0.2325 | 158 | 780 | 0.1785 | 0.0835 |
| >=2 | 0.1733 | 0.2085 | 375 | 1969 | 0.1785 | 0.1667 |
| >=3 | 0.1609 | 0.2057 | 217 | 1189 | 0.1785 | 0.7050 |
| >=4 | 0.1457 | 0.1975 | 144 | 845 | 0.1785 | 0.5595 |
| >=5 | 0.1399 | 0.1991 | 108 | 643 | 0.1785 | 0.5365 |
| >=6 | 0.1312 | 0.2000 | 77 | 471 | 0.1785 | 0.4340 |
| >=7 | 0.1202 | 0.1998 | 54 | 344 | 0.1785 | 0.3244 |
| >=8 | 0.1058 | 0.2140 | 29 | 188 | 0.1785 | 0.4460 |
| >=9 | 0.0998 | 0.2400 | 19 | 118 | 0.1785 | 0.7185 |
| >=10 | 0.0537 | 0.2490 | 7 | 54 | 0.1785 | 0.4760 |
| **K562** |  |  |  |  |  |  |
| ==2 | 0.1763 | 0.2271 | 200 | 996 | 0.1720 | 0.0186 |
| >=2 | 0.1767 | 0.2081 | 477 | 2484 | 0.1720 | 0.0091 |
| >=3 | 0.1667 | 0.2069 | 277 | 1488 | 0.1720 | 0.1491 |
| >=4 | 0.1491 | 0.1963 | 175 | 1019 | 0.1720 | 1.0000 |
| >=5 | 0.1354 | 0.1885 | 124 | 772 | 0.1720 | 0.4453 |
| >=6 | 0.1259 | 0.1878 | 86 | 555 | 0.1720 | 0.3113 |
| >=7 | 0.1040 | 0.1748 | 53 | 388 | 0.1720 | 0.0691 |
| >=8 | 0.0651 | 0.1451 | 24 | 240 | 0.1720 | 0.0026 |
| >=9 | 0.0393 | 0.1344 | 11 | 142 | 0.1720 | 0.0017 |
| >=10 | 0.0247 | 0.1656 | 5 | 67 | 0.1720 | 0.0343 |

## Notes

### Competing Interest Statement

The authors have declared no competing interest.

## References

1. Lupianez DG, Spielmann M, Mundlos S. Breaking TADs: How Alterations of Chromatin Domains Result in Disease. Trends Genet. 2016;32:225–37.

2. Lieberman-Aiden E, van Berkum NL, Williams L, Imakaev M, Ragoczy T, Telling A, Amit I, Lajoie BR, Sabo PJ, Dorschner MO, et al. Comprehensive mapping of long-range interactions reveals folding principles of the human genome. Science. 2009;326:289–93.

3. Rao SS, Huntley MH, Durand NC, Stamenova EK, Bochkov ID, Robinson JT, Sanborn AL, Machol I, Omer AD, Lander ES, Aiden EL. A 3D map of the human genome at kilobase resolution reveals principles of chromatin looping. Cell. 2014;159:1665–80.

4. Sexton T, Yaffe E, Kenigsberg E, Bantignies F, Leblanc B, Hoichman M, Parrinello H, Tanay A, Cavalli G. Three-Dimensional Folding and Functional Organization Principles of the Drosophila Genome. Cell. 2012;148:458–72.

5. Dixon JR, Selvaraj S, Yue F, Kim A, Li Y, Shen Y, Hu M, Liu JS, Ren B. Topological domains in mammalian genomes identified by analysis of chromatin interactions. Nature. 2012;485:376–80.

6. Larson AG, Elnatan D, Keenen MM, Trnka MJ, Johnston JB, Burlingame AL, Agard DA, Redding S, Narlikar GJ. Liquid droplet formation by HP1alpha suggests a role for phase separation in heterochromatin. Nature. 2017;547:236–40.

7. Strom AR, Emelyanov AV, Mir M, Fyodorov DV, Darzacq X, Karpen GH. Phase separation drives heterochromatin domain formation. Nature. 2017;547:241–45.

8. Sanborn AL, Rao SS, Huang SC, Durand NC, Huntley MH, Jewett AI, Bochkov ID, Chinnappan D, Cutkosky A, Li J, et al. Chromatin extrusion explains key features of loop and domain formation in wild-type and engineered genomes. Proc Natl Acad Sci U S A. 2015;112:E6456–65.

9. Fudenberg G, Imakaev M, Lu C, Goloborodko A, Abdennur N, Mirny LA. Formation of Chromosomal Domains by Loop Extrusion. Cell Rep. 2016;15:2038–49.

10. Nuebler J, Fudenberg G, Imakaev M, Abdennur N, Mirny LA. Chromatin organization by an interplay of loop extrusion and compartmental segregation. Proc Natl Acad Sci U S A. 2018;115:E6697–E706.

11. de Wit E. TADs as the caller calls them. J Mol Biol. 2019;10.1016/j.jmb.2019.09.026.

12. Rowley MJ, Nichols MH, Lyu X, Ando-Kuri M, Rivera ISM, Hermetz K, Wang P, Ruan Y, Corces VG. Evolutionarily Conserved Principles Predict 3D Chromatin Organization. Mol Cell. 2017;67:837–52 e7.

13. Beagan JA, Phillips-Cremins JE. On the existence and functionality of topologically associating domains. Nat Genet. 2020;52:8–16.

14. Paulsen J, Liyakat Ali TM, Nekrasov M, Delbarre E, Baudement MO, Kurscheid S, Tremethick D, Collas P. Long-range interactions between topologically associating domains shape the four-dimensional genome during differentiation. Nat Genet. 2019;51:835–43.

15. Davis CA, Hitz BC, Sloan CA, Chan ET, Davidson JM, Gabdank I, Hilton JA, Jain K, Baymuradov UK, Narayanan AK, et al. The Encyclopedia of DNA elements (ENCODE): data portal update. Nucleic Acids Res. 2018;46:D794–D801.

16. Filippova D, Patro R, Duggal G, Kingsford C. Identification of alternative topological domains in chromatin. Algorithms Mol Biol. 2014;9:14.

17. Paulsen J, Rodland EA, Holden L, Holden M, Hovig E. A statistical model of ChIA-PET data for accurate detection of chromatin 3D interactions. Nucleic Acids Res. 2014;42:e143.

18. Paulsen J, Sekelja M, Oldenburg AR, Barateau A, Briand N, Delbarre E, Shah A, Sørensen AL, Vigouroux C, Buendia B, Collas P. Chrom3D: three-dimensional genome modeling from Hi-C and lamin-genome contacts. Genome Biol. 2017;18:21.

19. Chen LL, Yang L. ALUternative Regulation for Gene Expression. Trends Cell Biol. 2017;27:480–90.

20. Elbarbary RA, Lucas BA, Maquat LE. Retrotransposons as regulators of gene expression. Science. 2016;351:aac7247.

21. Estecio MR, Gallegos J, Dekmezian M, Lu Y, Liang S, Issa JP. SINE retrotransposons cause epigenetic reprogramming of adjacent gene promoters. Mol Cancer Res. 2012;10:1332–42.

22. Lunyak VV, Prefontaine GG, Nunez E, Cramer T, Ju BG, Ohgi KA, Hutt K, Roy R, Garcia-Diaz A, Zhu X, et al. Developmentally regulated activation of a SINE B2 repeat as a domain boundary in organogenesis. Science. 2007;317:248–51.

23. Chow JC, Ciaudo C, Fazzari MJ, Mise N, Servant N, Glass JL, Attreed M, Avner P, Wutz A, Barillot E, et al. LINE-1 activity in facultative heterochromatin formation during X chromosome inactivation. Cell. 2010;141:956–69.

24. Rao SSP, Huang SC, Glenn St Hilaire B, Engreitz JM, Perez EM, Kieffer-Kwon KR, Sanborn AL, Johnstone SE, Bascom GD, Bochkov ID, et al. Cohesin Loss Eliminates All Loop Domains. Cell. 2017;171:305–20 e24.

25. An L, Yang T, Yang J, Nuebler J, Xiang G, Hardison RC, Li Q, Zhang Y. OnTAD: hierarchical domain structure reveals the divergence of activity among TADs and boundaries. Genome Biol. 2019;20:282.

26. Boettiger AN, Bintu B, Moffitt JR, Wang S, Beliveau BJ, Fudenberg G, Imakaev M, Mirny LA, Wu CT, Zhuang X. Super-resolution imaging reveals distinct chromatin folding for different epigenetic states. Nature. 2016;529:418–22.

27. Rowley MJ, Corces VG. Organizational principles of 3D genome architecture. Nat Rev Genet. 2018;19:789–800.

28. Eissenberg JC, Elgin SC. The HP1 protein family: getting a grip on chromatin. Curr Opin Genet Dev. 2000;10:204–10.

29. Servant N, Varoquaux N, Lajoie BR, Viara E, Chen CJ, Vert JP, Heard E, Dekker J, Barillot E. HiC-Pro: an optimized and flexible pipeline for Hi-C data processing. Genome Biol. 2015;16:259.

30. Hansen KD, Timp W, Bravo HC, Sabunciyan S, Langmead B, McDonald OG, Wen B, Wu H, Liu Y, Diep D, et al. Increased methylation variation in epigenetic domains across cancer types. Nat Genet. 2011;43:768–75.

31. Durand NC, Robinson JT, Shamim MS, Machol I, Mesirov JP, Lander ES, Aiden EL. Juicebox Provides a Visualization System for Hi-C Contact Maps with Unlimited Zoom. Cell Syst. 2016;3:99–101.

32. Csardi G, Nepusz T. The igraph software package for complex network research. InterJournal. 2006;Complex Systems:1695.

33. Bron C, Kerbosch J. Algorithm 457: finding all cliques of an undirected graph. Commun ACM. 1973;16:575–77.

34. Karolchik D, Hinrichs AS, Furey TS, Roskin KM, Sugnet CW, Haussler D, Kent WJ. The UCSC Table Browser data retrieval tool. Nucleic Acids Res. 2004;32:D493–D96.

35. Quinlan AR, Hall IM. BEDTools: a flexible suite of utilities for comparing genomic features. Bioinformatics. 2010;26:841–42.

36. Bruse N, Van Arensbergen J. GimmeMotifs: an analysis framework for transcription factor motif analysis. bioRxiv 474403. 2018;doi: https://doi.org/10.1101/474403.

